# Soil viral communities differed by management and over time in organic and conventional tomato fields

**DOI:** 10.1101/2023.05.03.539301

**Authors:** Jess W. Sorensen, Anneliek M. ter Horst, Laura A. Zinke, Joanne B. Emerson

## Abstract

Viruses are known contributors to biogeochemical cycling in diverse habitats, but viral community studies in soil are relatively rare. Although prior work has suggested spatial structuring as a primary driver of local soil viral community ecological patterns, here we show that agricultural management can significantly impact soil viral community composition. We generated 18 soil viromes and 24 total metagenomes from six plots, three under organic and three under conventional management in Davis, CA, USA. Despite the significant impact of management on viral community structure and soil physicochemistry, approximately 44% of the detected viral ‘species’ (vOTUs) were present in soils from both management practices. These common vOTUs regularly comprised >65% of the viral community by relative abundance. Many (56%) of the vOTUs were detected both during the tomato growing season and post-harvest, indicating persistence through time. Together, these results indicate habitat-specific differences in viral community composition, yet relative stability and persistence of viral communities within agricultural soils, in contrast to their recently observed dynamics in natural soils.

## Introduction

There is a growing body of work indicating the importance of viruses in biogeochemical cycling (1). While technical challenges have made accessing soil viruses difficult, methodological improvements have revealed an incredible diversity of viruses in soil and their potential roles in biogeochemical cycles (2–4). For example, soil viruses likely influence methane cycling through infection of methanogens and methanotrophs (5–7), and they can impact soil carbon and nitrogen cycling in myriad ways, including via expression of viral auxiliary metabolic genes (3, 8–10). Recent studies have shown differences between agricultural and more natural soils in their endemic viral communities (11–13), as well as the potential for cropping rotations to influence soil viral community composition (14). However, the influence of other agricultural management practices, such as organic and conventional management, on soil viruses is still relatively unknown.

Here we report the impacts of two different agricultural management practices on soil viral communities associated with tomato cropping in six one-acre plots at the UC Davis Russell Ranch Sustainable Agriculture Facility’s Century Experiment in Davis, California, USA (Fig. S1) (15). Prior to sample collection in 2018, management practices had been underway since 1994, with all plots in a tomato-corn rotation with a primary crop of tomato or corn in alternate years. Three plots had been organically managed, receiving poultry manure compost (9 tonnes per hectare) every fall post-harvest (November) and planted with a cover crop mix of purple lana vetch, bell bean, and clover during the winter months (November to March). The remaining three plots had been conventionally managed, receiving 156 kg per hectare of mineral fertilizer (urea ammonium nitrate solution) at 3-4 times throughout the tomato (or corn) growing season, and left unplanted (fallow) during the winter months. All six plots were tilled once per year in the spring and received water via subsurface drip irrigation throughout the summer. We prepared and sequenced soil viromes (post-0.22 μm‘viral size fraction’ metagenomes) and total soil metagenomes from the six plots during the tomato growing season (July) and post-harvest (October) to assess the influence of the two management practices on soil viral communities in comparison to prokaryotic communities.

## Results and Discussion

From the three organically managed and three conventionally managed tomato-corn rotation plots, we generated a total of 18 viromes (one per plot in July, two per plot in October) and 24 total soil metagenomes (two per plot at both time points). A full description of sampling procedures, virome preparation, nucleic acid extraction, sequencing, and analysis can be found online (Supplementary Methods). A total of 7,944 viral operational taxonomic units (vOTUs) >10 kbp was detected in the viromes, with a median of 1,244 vOTUs per virome (Table S1), and 278,959 contigs >2 kbp and 750,900 reads identified as 16S rRNA genes were recovered from the total metagenomes.

We were interested in whether agricultural management and/or time period significantly impacted soil viral and/or prokaryotic community composition. Both management and sampling date were significantly correlated with viral community structure, though management had a stronger effect (Fig. 1A, PERMANOVA p = 0.0002 for both management [pseudo-F = 4.884] and date [pseudo-f = 3.8915]). The impacts of management and sampling date on the bacterial and archaeal communities were less significant. Specifically, microbial communities measured through read-mapping-based contig relative abundances were significantly correlated with management and time (Fig. 1B, PERMANOVA p=0.0001 [pseudo-F = 3.477] and p=0.0001 [pseudo-F = 4.669], respectively), but the strength of these correlations was noticeably lower for the 16S rRNA gene fragments recovered from these metagenomes (Fig. S2, PERMANOVA p=0.023 [pseudo-F = 2.410], p=0.0004 [pseudo-F = 4.669, respectively).

**Figure 1.**
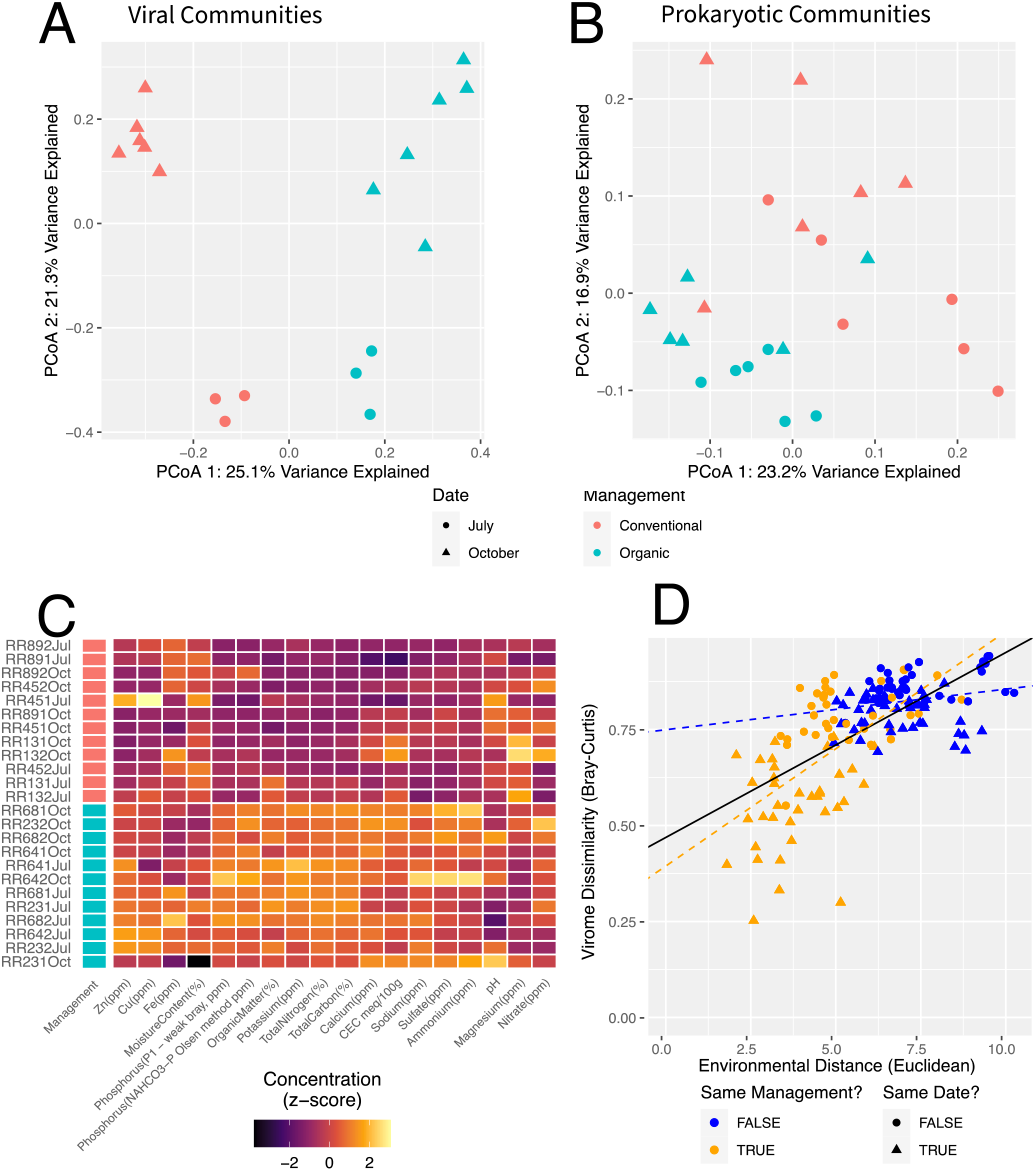
Viral and microbial community structure differed by management and time of year. (A and B) Principal Coordinates Analysis (PCoA) plots of Bray-Curtis dissimilarities between (A) viral and (B) prokaryotic communities. Viral community composition was derived from read mapping to vOTUs for relative abundances, and prokaryotic community composition was derived from read mapping to a dereplicated set of metagenomic contigs > 2 kbp. Each point is a single community, with color mapped to agricultural management and shape mapped to date of sample collection. (C) Heatmap showing z-scores of the relative concentrations or values (for pH) of 18 soil physicochemical properties across our 24 samples. Each row represents a single sample with a unique sample ID (left). Management is colored according to the legend below panels A and B (pink for conventional, blue for organic). (D) Viral community Bray-Curtis dissimilarities plotted against Euclidean environmental distances calculated from the 18 physicochemical properties shown in panel C. Each point represents a pairwise sample-to-sample comparison. Points are colored based on whether the pair of compared samples belonged to the same agricultural management (organic-organic or conventional-conventional, TRUE) or different management practices (organic-conventional, FALSE), and the shape of each point indicates whether the compared samples were from the same collection date (TRUE) or different dates (FALSE). Lines represent the linear regressions for: the full dataset (all points, black), all pairwise comparisons within the same management (orange), and all pairwise comparisons between managements (blue).

To investigate what aspects of the agricultural management practices might be influencing soil viral community composition, we next considered the chemistry of the soils. Multiple soil physicochemical properties were significantly different between management practices, including organic matter, total carbon, and total nitrogen (Figure 1C, Table S2). In particular, soil organic matter concentrations were significantly higher in the organically managed soils, suggesting that the annual application of poultry manure compost and winter cover crop indeed altered the soil chemistry in these plots. To assess whether soil physicochemical properties in aggregate could explain the observed differences in viral community structure, we calculated pairwise Euclidean distances between samples, using the z-scored soil chemistry profiles, and compared them to viral community Bray-Curtis dissimilarities. Environmental distances were significantly correlated with soil viral community dissimilarities (Figure 1D, Spearman’s rho = 0.653, p <2.2 ×10^−16^), with less similar communities over increasing environmental distances. This relationship held for comparisons between samples belonging to the same management (organic-organic or conventional-conventional, Figure 1D, Spearman’s rho=0.576, p = 2.693 × 10^−14^). However, when restricting our analyses only to comparisons between samples belonging to different management practices, the relationship between environmental distances and Bray-Curtis dissimilarities was much weaker (had a flatter slope), though was still significant (Spearman’s rho=0.204, p=0.009). This reduction in significance may indicate the presence of viral community alternative stable states (16) across management practices. In other words, environmental dissimilarity may correlate with viral community dissimilarity up to a threshold at which the environmental differences become so great as to select for fundamentally different communities. These same patterns were observed in the total prokaryotic communities as well, though the relationship between community dissimilarity and environmental distance was weaker (Spearman’s rho = 0.384, p <2.2 ×10^−16^, Figure S3). This weaker relationship in prokaryotic communities is indicative of slower community turnover, and/or it could reflect the presence of dormant microbes and relic DNA (17–19) in our prokaryotic community measurements, perhaps masking patterns in the active microbiome.

We next wanted to identify changes at the viral ‘species’ (vOTU) level underlying the observed differences in community composition according to management. Conventionally and organically managed plots showed no significant difference in viral richness (the number of vOTUs detected, Fig. S4, Kruskal-Wallis p>0.05). In total, 3,516 vOTUs (or 44%) were detected in plots from both management practices, while 2,699 (34%) and 1,729 (22%) were detected only in conventionally or organically managed soils, respectively (Fig. 2A). This relatively high degree of overlap in vOTUs throughout the farm runs contrary to observations in natural systems, in which the vast majority of vOTUs tend to be spatially restricted (e.g., only recovered in single habitats or on one end of a field, if not in single samples (19, 20)). Not only was a large proportion of vOTUs shared between management practices here, these shared vOTUs regularly comprised >65% of the relative abundance in these viral communities (Fig. 2B). This pattern of persistent viral populations in soil also extended through time. Of the 7,944 total vOTUs, 4,448 (56%) were observed in both July and October (Fig S5A), and these vOTUs comprised on average 88% of the relative abundance of a given viral community (Fig S5B). Taken together, these data suggest the persistence of a core set of agricultural soil viral ‘species’ over space and time, despite differences in management.

**Figure 2.**
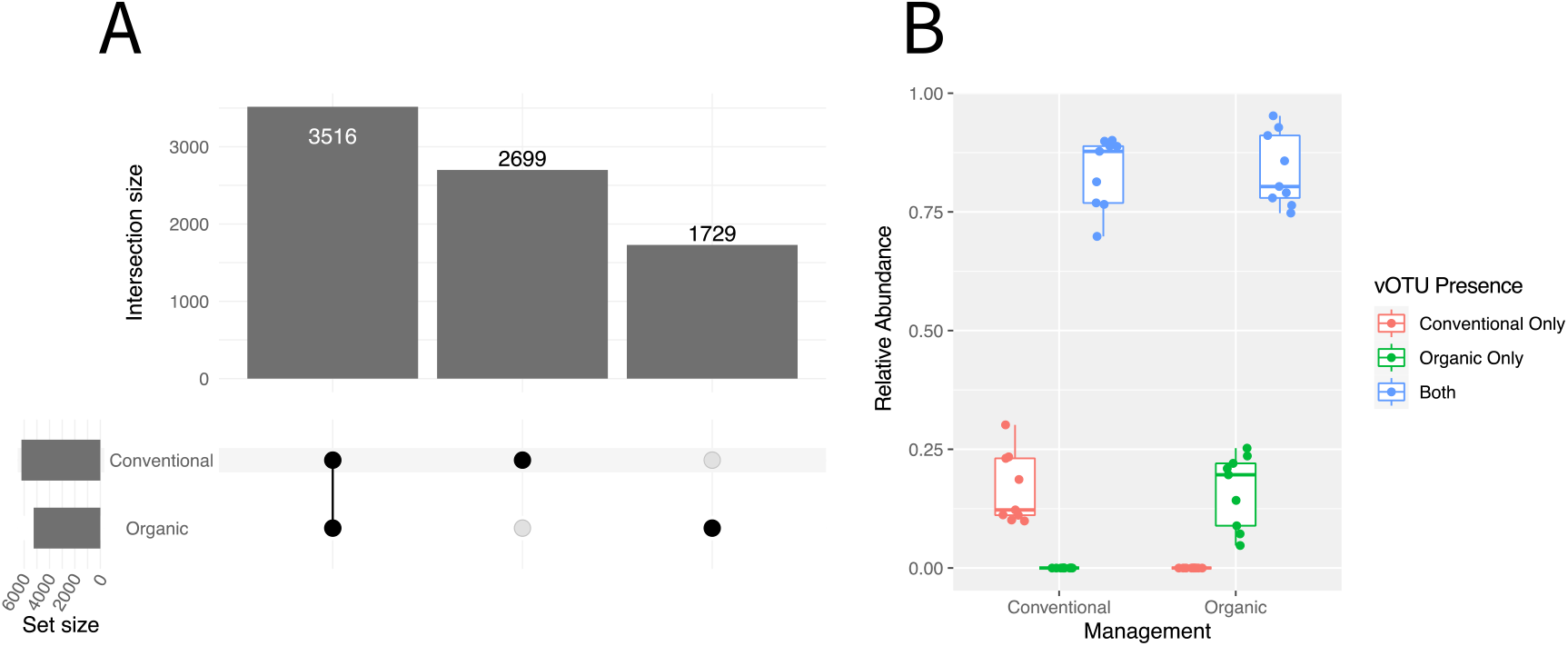
Most vOTUs were detected in soils from both agricultural management practices. (A) An upset plot showing the number of vOTUs (intersection size) shared between (connected black dots) and unique to (single black dots) each of the two agricultural management practices. Set size on the left indicates the total number of vOTUs detected within each management practice. (B) Summed relative abundances of vOTUs unique to each management practice (green or pink) or shared across management practices (blue). Each point is one sample. Lines in each box represent the median, boxes extend to the 25^th^ and 75^th^ percentile, whiskers extend to the furthest point within 1.5 times the inter-quartile range, and points beyond whiskers are outliers.

Unlike recent studies of soil viral communities (20, 21), the results here present a picture of a relatively persistent set of soil viral populations, with differences in community composition attributable to the soil physicochemical environment. Recent studies have repeatedly detected spatial organization of viral communities (19, 21, 22), including at another tomato research field site in Davis, CA (22), but there was no observation of such spatial structuring at the scales examined in this study. Instead, results align with another growing body of work that highlights the importance of habitat and land use in structuring soil viral communities (14, 20, 23, 24). For example, in a recent study of agricultural and early successional soils (soils dominated by annuals and grasses), the agricultural soil viral communities were the most stable over the growing season (12), consistent with the observation here of a relatively large proportion of persistent agricultural soil vOTUs over time. Agricultural soils are different from most other soils in ways that are somewhat obvious, but that, as the current study has revealed, likely have substantial impacts on viral ecological dynamics. For example, these soils are commonly mixed through processes such as tilling, they share the same crop plant, and they are managed with relatively homogeneous inputs, such as fertilizer (albeit in different forms by management here) and irrigation, which together lead to more ecosystem homogeneity than would be observed in natural soils (12, 13). It thus follows that the potential influence of spatial structuring would be minimized in agricultural soils that experience homogenization and repeated disruptions of soil structure.

## Data Availability Statement

Raw sequencing reads from viromes and metagenomes, as well as vOTU sequences, are available under BioProject accession PRJNA767554.

## Acknowledgements

The research described herein was primarily supported by USDA NIFA AFRI award # 2021-67013-34815-0 (grant to JBE). Additional support was provided by USDA NIFA Hatch # CA-D-PPA-2464-H. JWS was supported by USDA National Institute of Food and Agriculture award # 2021-67034-35002. Thanks to Sara Geonczy and Rose Bolle for assistance with sample collection, and thanks to Israel Herrera and Nicole Tautges for coordinating access to the Russell Ranch Sustainable Agriculture Facility and its historical management data.

## References

1. Roux S, Emerson JB. 2022. Diversity in the soil virosphere: to infinity and beyond? Trends Microbiol 30:1025–1035.

2. Godinez I, Mcdermott JE, Hofmockel KS, Jansson K. 2021. DNA Viral Diversity, Abundance, and Functional Potential Vary across Grassland Soils with a Range of Historical Moisture Regimes. MBio 12:e02595–21.

3. Wu R, Smith CA, Buchko GW, Blaby IK, Paez-Espino D, Kyrpides NC, Yoshikuni Y, McDermott JE, Hofmockel KS, Cort JR, Jansson JK. 2022. Structural characterization of a soil viral auxiliary metabolic gene product – a functional chitosanase. Nat Commun 13:1–14.

4. Zheng X, Jahn MT, Sun M, Friman VP, Balcazar JL, Wang J, Shi Y, Gong X, Hu F, Zhu YG. 2022. Organochlorine contamination enriches virus-encoded metabolism and pesticide degradation associated auxiliary genes in soil microbiomes. ISME J 16:1397–1408.

5. Emerson JB, Roux S, Brum JR, Bolduc B, Woodcroft BJ, Jang H Bin, Singleton CM, Solden LM, Naas AE, Boyd JA, Hodgkins SB, Wilson RM, Trubl G, Li C, Frolking S, Pope PB, Wrighton KC, Crill PM, Chanton JP, Saleska SR, Tyson GW, Rich VI, Sullivan MB. 2018. Host-linked soil viral ecology along a permafrost thaw gradient. Nat Microbiol 3:870–880.

6. Lee S, Sieradzki ET, Nicolas AM, Walker RL, Firestone MK, Hazard C, Nicol GW. 2021. Methane-derived carbon flows into host-virus networks at different trophic levels in soil. Proc Natl Acad Sci U S A 118:1–8.

7. Trubl G, Jang H Bin, Roux S, Emerson JB, Solonenko N, Vik DR, Solden L, Ellenbogen J, Runyon AT, Bolduc B, Woodcroft BJ, Saleska SR, Tyson GW, Wrighton KC, Sullivan MB, Rich VI. 2018. Soil viruses are underexplored players in ecosystem carbon processing. mSystems 3:1–21.

8. Albright MBN, Gallegos-Graves LV, Feeser KL, Montoya K, Emerson JB, Shakya M, Dunbar J. 2022. Experimental evidence for the impact of soil viruses on carbon cycling during surface plant litter decomposition. ISME Commun 2:1–8.

9. Braga LPP, Spor A, Kot W, Breuil MC, Hansen LH, Setubal JC, Philippot L. 2020. Impact of phages on soil bacterial communities and nitrogen availability under different assembly scenarios. Microbiome 8:1–14.

10. Starr EP, Shi S, Blazewicz SJ, Koch BJ, Probst AJ, Hungate BA, Pett-Ridge J, Firestone MK, Banfield JF. 2021. Stable-Isotope-Informed, Genome-Resolved Metagenomics Uncovers Potential Cross-Kingdom Interactions in Rhizosphere Soil.mSphere 6.

11. Liao H, Li H, Duan CS, Zhou XY, Luo QP, An XL, Zhu YG, Su JQ. 2022. Response of soil viral communities to land use changes. Nat Commun 13.

12. Roy K, Ghosh D, DeBruyn JM, Dasgupta T, Wommack KE, Liang X, Wagner RE, Radosevich M. 2020. Temporal Dynamics of Soil Virus and Bacterial Populations in Agricultural and Early Plant Successional Soils. Front Microbiol 11.

13. Cornell CR, Zhang Y, Van Nostrand JD, Wagle P, Xiao X, Zhou J. 2021. Temporal Changes of Virus-Like Particle Abundance and Metagenomic Comparison of Viral Communities in Cropland and Prairie Soils. mSphere 6.

14. Muscatt G, Hilton S, Raguideau S, Teakle G, Lidbury IDEA, Wellington EMH, Quince C, Millard A, Bending GD, Jameson E. 2022. Crop management shapes the diversity and activity of DNA and RNA viruses in the rhizosphere. Microbiome 10:1–16.

15. Wolf KM, Torbert EE, Bryant D, Burger M, Denison RF, Herrera I, Hopmans J, Horwath W, Kaffka S, Kong AYY, Norris RF, Six J, Tomich TP, Scow KM. 2018. The century experiment: the first twenty years of UC Davis’ Mediterranean agroecological experiment. Ecology 99:503.

16. Mushet DM, McKenna OP, McLean KI. 2020. Alternative stable states in inherently unstable systems. Ecol Evol 10:843–850.

17. Lennon JT, Jones SE. 2011. Microbial seed banks: the ecological and evolutionary implications of dormancy. Nat Rev Microbiol 9:119–30.

18. Locey KJ, Muscarella ME, Larsen ML, Bray SR, Jones SE, Lennon JT. 2020. Dormancy dampens the microbial distance-decay relationship. Philos Trans R Soc B Biol Sci 375.

19. Santos-Medellín C, Estera-Molina K, Yuan M, Pett-Ridge J, Firestone MK, Emerson JB. 2022. Spatial turnover of soil viral populations and genotypes overlain by cohesive responses to moisture in grasslands. Proc Natl Acad Sci U S A 119:1–11.

20. Durham DM, Sieradzki ET, ter Horst AM, Santos-Medellín C, Bess CWA, Geonczy SE, Emerson JB. 2022. Substantial differences in soil viral community composition within and among four Northern California habitats. ISME Commun 2:1–5.

21. Horst AM, Santos-medellín C, Sorensen JW, Zinke LA, Wilson RM, Johnston ER, Trubl G, Pett-ridge J, Blazewicz SJ, Hanson PJ, Chanton JP, Schadt CW, Kostka JE, Emerson JB. 2021. Minnesota peat viromes reveal terrestrial and aquatic niche partitioning for local and global viral populations. Microbiome 9:1–19.

22. Santos-Medellin C, Zinke LA, ter Horst AM, Gelardi DL, Parikh SJ, Emerson JB. 2021. Viromes outperform total metagenomes in revealing the spatiotemporal patterns of agricultural soil viral communities. ISME J https://doi.org/10.1101/2020.08.06.237214.

23. Lee S, Sorensen JW, Walker RL, Emerson JB, Nicol GW, Hazard C. 2022. Soil pH influences the structure of virus communities at local and global scales. Soil Biol Biochem 166:108569.

24. Hillary LS, Adriaenssens EM, Jones DL, McDonald JE. 2022. RNA-viromics reveals diverse communities of soil RNA viruses with the potential to affect grassland ecosystems across multiple trophic levels. ISME Commun 2:1–10.

