## Supplemental_Methods_Figures for "Soil viral communities differed by management and over time in organic and conventional tomato fields"

*Study design, sampling procedure, and soil chemistry*

This study focused on three conventionally managed and three organically managed one-acre (64m x 64m) tomato-corn rotation plots at the UC Davis Russell Ranch Sustainable Agricultural Facility in Davis, CA, USA. In July and October of 2018 (shortly before and approximately one month after tomato harvest), we sampled three conventionally managed (CMT) and three organically managed (OMT) plots. All plots were irrigated through subsurface drip irrigation systems and were growing tomatoes during the 2018 growing season. Poultry manure compost was applied to the organically managed fields post-harvest each year (November 2017), while conventionally managed plots were fertigated (i.e., fertilizer was applied via drip irrigation) with mineral fertilizer (urea ammonium nitrate) 3-4 times during the tomato growing season. Additionally, conventionally managed fields were left unplanted (fallow) during the fall and winter, while organically managed plots were planted with a cover crop mixture of bell bean, lana vetch, and clover.

Three five cm diameter cores of the top 15 cm of soil were collected per sample, and two samples per plot were collected per time point, resulting in a total of 24 samples. The two samples per plot (per time point) were collected on opposite ends of the plot 32 meters apart and 12 meters away from edges to avoid edge effects (Table S1). Individual soil samples (triplicate cores) were passed through an 8 mm sieve for homogenization. Soils were sent to A&L Western Agricultural Labs in Modesto, CA for soil chemistry profiles, while nitrate, ammonium, total nitrogen, and total carbon were measured at the UC Davis Analytical Lab. Gravimetric soil moisture was measured via measuring out 10g fresh soil and leaving it at room temperature, massing daily until completely dry (i.e. mass stabilized). [add method].

*Soil Viromics, DNA extraction, Library Construction, and Sequencing*

Soil viromes were extracted following a previously published protocol (1), with slight modifications (2). Extractions were performed on 100 g total of frozen soil (stored at -80 C between sample collection and processing). Briefly, for each sample, four 50 mL conical tubes were filled with 25 g frozen soil and 37.5 mL 0.02 μm-filtered AKC' buffer (10 g Potassium Citrate, 1.44 g $Na_2HPO_4$, 0.24 g $KH_2PO_4$, 36.97 g $MgSO_4$, 10% PBS, per liter of MilliQ water). Tubes were shaken at 400 rpm for 15 minutes on an orbital shaker, before being vortexed for three minutes and centrifuged at 4,700 g for 15 minutes. Supernatants were then filtered through a 0.22 μm filter to remove most cells. The four supernatants per sample were then reduced to two by combining two supernatants into a single 70 mL ultracentrifuge tube. Supernatants were then spun at 32,000 g for 3 hours at 4°C using an Optima LE-80K ultracentrifuge (Beckman-Coulter Life Sciences, Indianapolis IN). Resulting supernatants were decanted, and the two pellets per sample were resuspended and combined in a total of 200 μL of ultrapure water.

Each viral suspension was treated with 30 units of RQ1 RNase-free DNase (Promega Corporation, Madison, WI USA) for two hours at room temperature. Reactions were quenched through the addition of 1μl of stop solution and incubation at 65°C for 10 minutes. DNA was extracted from the DNase-treated viral suspensions using the DNeasy PowerSoil Kit (Qiagen, Hilden, Germany) with slight modifications. The bead beating step was replaced with a 10-minute 70°C incubation, a 5-second vortex, and another 5-minute, 70°C incubation. Extracted DNA was quantified using the 1X High Sensitivity DNA assay on a Qubit 4 Fluorometer (ThermoFischer Scientific, Inc. Watham, MA USA). Samples with DNA below detection limits were not submitted for sequencing, resulting in only one virome per plot for the July samples.

For total soil metagenomes, DNA was extracted from 0.5 g of soil per sample, using the DNeasy PowerSoil Kit (Qiagen, Hilden, Germany), according to the manufacturer's instructions.

For both viromes and total metagenomes, library construction and sequencing were performed by the DNA Technologies & Expression Analysis Core Laboratory at UC Davis (Davis, CA). Libraries were constructed with the DNA KAPA HyperPrep kit (Kapa Biosystems-Roche, Basel, Switzerland). Paired-end 150 bp sequencing was performed on an Illumina HiSeq4000 (Illumina) to a requested depth of 40,000,000 read pairs per sample (actual sequencing depths in Table S1).

*Sequence quality checking, assembly, viral contig identification, and analysis*

Adapters, primers, and PhiX sequences were removed from the libraries using BBDUK (3). Reads were then quality trimmed using Trimmomatic (4), using a quality cutoff score of 30 in a 4-base sliding window. Reads shorter than 50 bp after trimming were removed. Quality filtered and trimmed reads from each sample were then individually *de novo* assembled using MEGAHIT (5) with a minimum contig length of 10 kb for viromes and 2 kb for total soil metagenomes. Resulting contigs within each library type (viromes and total soil metagenomes) were then clustered using PSI-CD-HIT (6) at a global identity threshold of 0.95.

Viral contigs were identified from the viromes using VirSorter (7) in virome mode and DeepVirFinder (8). For VirSorter, only viruses predicted in quality categories 1, 2, 4, and 5 were retained. Likewise, only DeepVirFinder predicted viruses with scores >0.9 and p-values <0.05 were retained. Reads were then mapped at 90% average nucleotide identity using bbmap (3) to this set of contigs and to our in-house database of phages, PIGEONv1.0 (9), to identify as many putative viral contigs as possible. Only viral contigs with >75% breadth (>75% bases covered at least 1x) via read mapping were considered in downstream analyses. We used the same mapping identity and base coverage thresholds for detection of microbial contigs assembled from total soil metagenomes. Partial 16S rRNA gene sequences were recovered from total metagenomes using SortMeRNA(10).

*Ecological Analyses*

Alpha and beta diversity analyses were performed in R (11), using the vegan package (12). All plots were constructed using ggplot2 (13). Significant differences in viral community structure attributable to experimental factors (management and time) were assessed with PERMANOVA, using the adonis() function from the vegan package (12). Correlations between viral community dissimilarity and different soil chemical profiles were assessed with a Mantel test using the mantel() function from the vegan package (12). Linear regressions seen in Figure 1D were performed using the lm() function in R.


1. Trubl G, Solonenko N, Chittick L, Solonenko SA, Rich VI, Sullivan MB. 2016. Optimization of viral resuspension methods for carbon-rich soils along a permafrost thaw gradient. PeerJ 4:1–24.
2. Santos-Medellin C, Zinke LA, ter Horst AM, Gelardi DL, Parikh SJ, Emerson JB. 2021. Viromes outperform total metagenomes in revealing the spatiotemporal patterns of agricultural soil viral communities. ISME J https://doi.org/10.1101/2020.08.06.237214.
3. Bushnell B. BBTools. https://sourceforge.net/projects/bbmap/.
4. Bolger AM, Lohse M, Usadel B. 2014. Trimmomatic: A flexible trimmer for Illumina sequence data. Bioinformatics 30:2114–2120.
5. Li D, Liu C-M, Luo R, Sadakane K, Lam T-W. 2015. MEGAHIT: an ultra-fast single-node solution for large and complex metagenomics assembly via succinct de Bruijn graph. Bioinformatics 31:1674–1676.
6. Fu L, Niu B, Zhu Z, Wu S, Li W. 2012. CD-HIT: Accelerated for clustering the next-generation sequencing data. Bioinformatics 28:3150–3152.
7. Roux S, Enault F, Hurwitz BL, Sullivan MB. 2015. VirSorter: Mining viral signal from microbial genomic data. PeerJ 2015:1–20.
8. Ren J, Song K, Deng C, Ahlgren NA, Fuhrman JA, Li Y, Xie X, Poplin R, Sun F. 2020. Identifying

viruses from metagenomic data using deep learning. Quant Biol 8:64–77.

9. Horst AM, Santos-medellín C, Sorensen JW, Zinke LA, Wilson RM, Johnston ER, Trubl G, Pett-ridge J, Blazewicz SJ, Hanson PJ, Chanton JP, Schadt CW, Kostka JE, Emerson JB. 2021. Minnesota peat viromes reveal terrestrial and aquatic niche partitioning for local and global viral populations. Microbiome 9:1–19.

10. Kopylova E, Noé L, Touzet H. 2012. SortMeRNA: Fast and accurate filtering of ribosomal RNAs in metatranscriptomic data. Bioinformatics 28:3211–3217.

11. R Core Team. 2017. R: A Language and Environment for Statistical Computing. Vienna, Austria.

12. Oksanen J, Blanchet FG, Friendly M, Kindt R, Legendre P, McGlinn D, Minchin PR, O'Hara RB, Simpson GL, Solymos P, Stevens MHH, Szoecs E, Wagner H. 2017. vegan: Community Ecology Package.

13. Wickham H. 2009. ggplot2: Elegant Graphics for Data Analysis. Springer-Verlag New York.

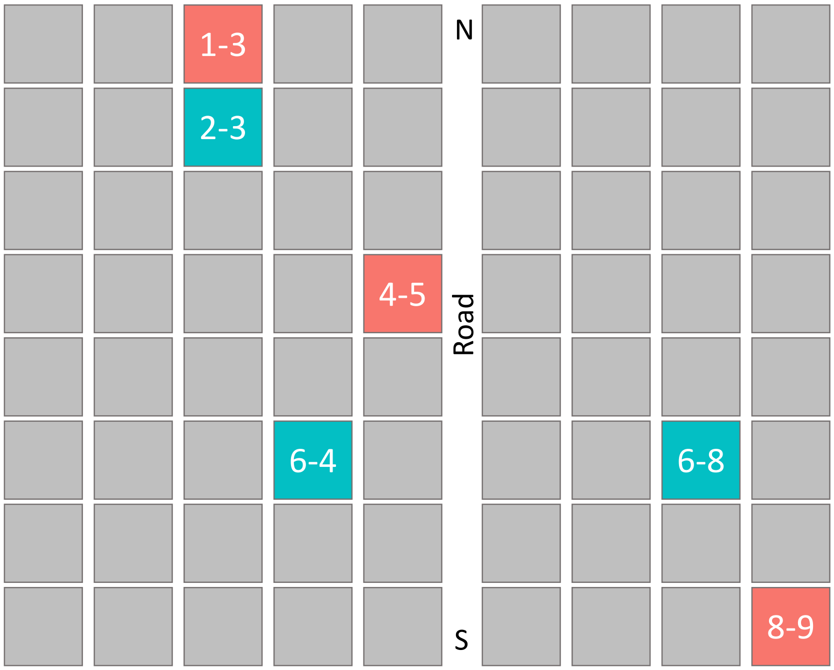


**Figure S1.** Map of the UC Davis Russell Ranch Sustainable Agriculture Facility's Century Experiment. The seventy-two one-acre plot design of the Russell Ranch Century Experiment is shown (most plots, colored grey, were not sampled for this study). The six sampled Tomato-Corn rotation plots in this study are highlighted, with organically managed plots shown in blue and conventionally managed shown in red.

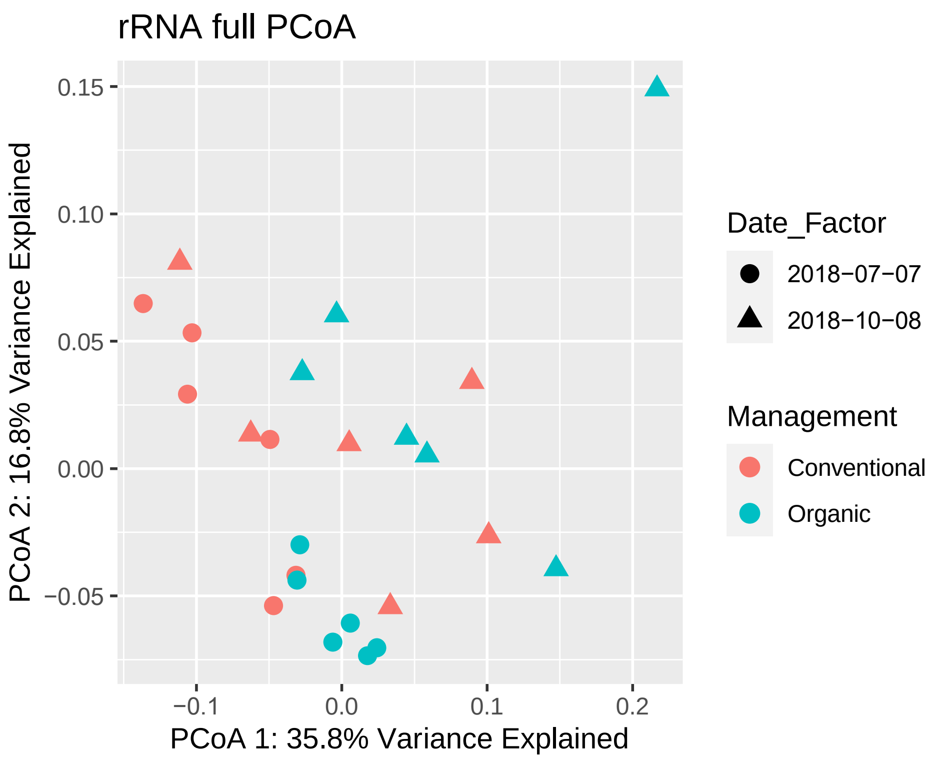


**Figure S2.** Microbial community structure across 24 soil samples, as measured by partial 16S rRNA gene fragments recovered from total metagenomes using SortMeRNA. This set of samples matches the set with paired viromes (Figure 1A) and is a subset of the 24 samples depicted in Figure 1B. Each point represents the microbial community associated with a particular sample. The color of each point corresponds to the agricultural management practice for the plot, while shape corresponds to the sampling date. The graph was constructed, using Principal Coordinates Analysis of the Bray-Curtis dissimilarities between microbial communities.

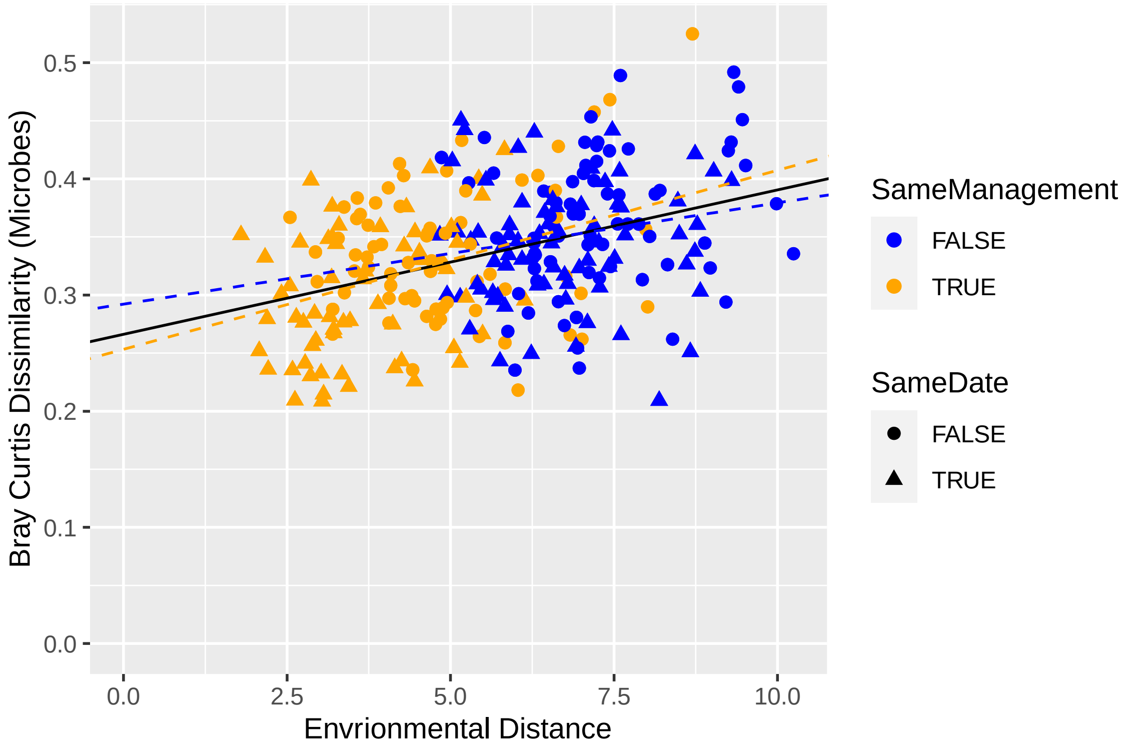


**Figures S3.** Relationship between microbial community structure and environmental distance. Prokaryotic community Bray-Curtis dissimilarities plotted against Euclidean environmental distances calculated from the 18 physicochemical properties shown in Fig. 1C. Each point represents a pairwise sample-to-sample comparison. Points are colored based on whether the pair of compared samples belonged to the same agricultural management (organic-organic or conventional-conventional, TRUE) or different management practices (organic-conventional, FALSE), and the shape of each point indicates whether the compared samples were from the same collection date (TRUE) or different dates (FALSE). Lines represent the linear regressions for: the full dataset (all points, black), all pairwise comparisons within the same management (orange), and all pairwise comparisons between managements (blue).

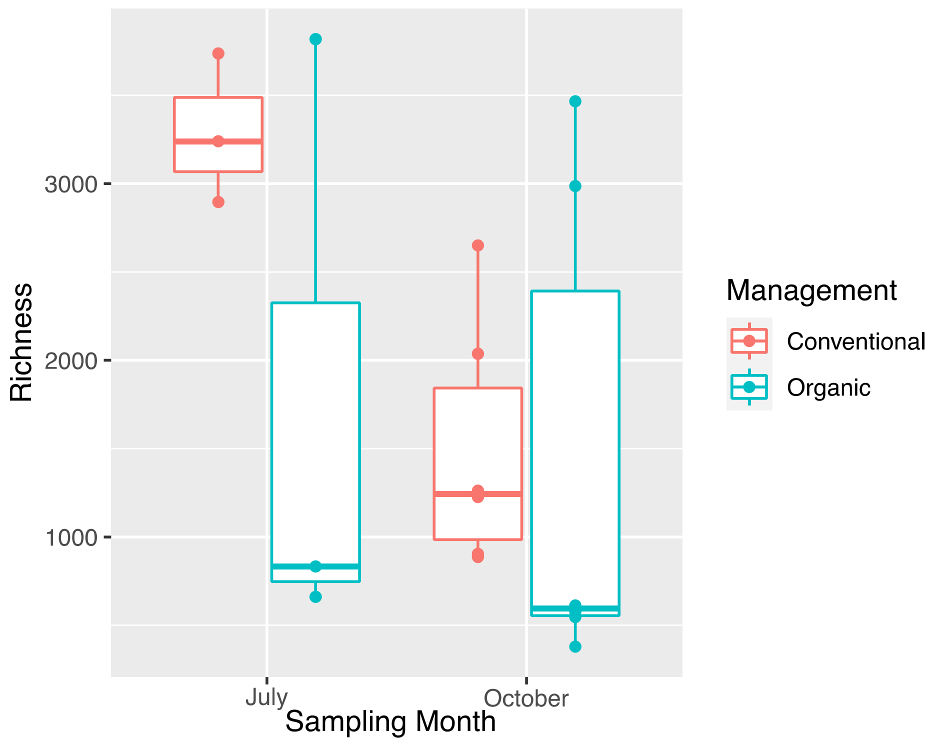


**Figure S4.** Viral community richness did not differ significantly between management practices. Richness is defined here as the number of vOTUs with ≥ 75% of their sequence length covered by at least one read and an average coverage depth ≥ 1. Each point represents the richness of a single soil virome, with points colored based on management. Lines in each box represent the median, boxes extend to the 25$^{th}$ and 75$^{th}$ percentile, whiskers extend to the furthest point within 1.5 times the inter-quartile range, and points beyond whiskers are outliers.

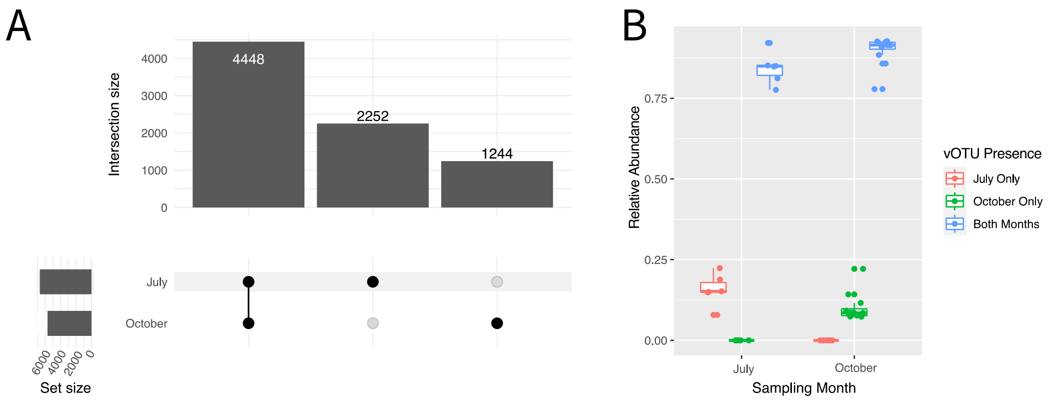


**Figure S5.** vOTUs persist in soils through time. (A) An upset plot showing the number of vOTUs (intersection size) shared between (connected black dots) and unique to (single black dots) each of the two sampling dates. Set size on the left indicates the total number of vOTUs detected within each management practice. (B) Summed relative abundances of vOTUs unique to each sampling date (green or pink) or shared across sampling dates (blue). Each point is the summed relative abundance of a set of vOTUs in one sample. Lines in each box represent the median, boxes extend to the 25$^{th}$ and 75$^{th}$ percentile, whiskers extend to the furthest point within 1.5 times the inter-quartile range, and points beyond whiskers are outliers.
